# Formic acid impairs α-synuclein seeding activity in human and mouse brains

**DOI:** 10.1101/2025.07.14.664807

**Authors:** Saumya Digraskar, George Dobbins, Piyali Das, Ryan Bash, Abigail Songer, Jaewon Huh, Kasandra Scholz, Musaab Khan, Matthew S Goldberg, Marla Gearing, Laura Volpicelli-Daley, Shu Chen, Patricia Aguilar-Calvo

## Abstract

α-Synuclein (α-syn) is an abundant monomeric protein that can aggregate into fibrils and form neuropathological inclusions in the brains of patients with synucleinopathies. New evidence suggests that the mouse-human transmission barrier of α-syn is lower than previously reported, emphasizing the need for improved biosafety procedures when working with α-syn aggregates. Histopathology of α-syn-infected brain represents a significant potential source of occupational exposure, and current methods for tissue fixation do not inactivate the ability of pathologic α-syn to seed the conversion of endogenous, monomeric α-syn into fibrils. In this study, we tested whether 96% formic acid treatment could reduce the seeding activity of α-syn aggregates in paraformaldehyde-fixed brain samples from dementia with Lewy bodies (DLB) patients and α-syn pre-formed fibrils (PFF)-injected mouse brains. Using real-time quaking-induced conversion (RT-QuIC), we found that formic acid treatment reduced α-syn seeding dose (picograms of α-syn seeds per ml of brain homogenate) in DLB and mouse brain by 6 and 8 logarithms, respectively. RT-QuIC reactions seeded with formic acid-treated brain homogenates showed significantly longer lag phase, and decreased total thioflavin T fluorescence compared to untreated samples, indicating that formic acid treatment impairs the ability of pathological α-syn to seed monomeric α-syn. Importantly, the α-syn pathologic features and the immunostaining quality were preserved in formic acid-treated tissues. Our results demonstrate that formic acid treatment is a quick and efficient procedure for reducing α-syn seeding activity in fixed brain samples, thereby lowering the risk of accidental exposure in laboratories without compromising the quality of histopathological analysis.

**Summary:** Formic acid treatment drastically reduces α -synuclein seeding activity in fixed human and mouse brain samples while preserving histopathological quality.

## Introduction

Synucleinopathies such as Parkinson’s disease (PD) and dementia with Lewy bodies (DLB) are caused by the misfolding and aggregation of α-synuclein (α-syn) protein in the central nervous system. α-Syn aggregates induce the conversion of endogenous α-syn into neuronal inclusions, called Lewy bodies (LB) and Lewy Neurites (LN), immunoreactive for α-syn phosphorylated at serine 129 (p-α-syn), a relatively selective marker of α-syn pathology in human postmortem brains (*1*) and animal models (*2*). Although the precise mechanisms of α-syn aggregation are not fully understood, it is thought to occur through a seeded or templated polymerization process. This self-replication process, often compared to the prion-like propagation mechanisms, advances α-syn pathology to further brain areas (*3*).

The intracranial injection of mice with fragmented pre-formed fibrils (PFFs) of recombinant α-syn reliably induces the conversion of endogenous α-syn into phosphorylated and filament enriched α-syn deposits, as well as synaptic dysfunction, altered neuronal excitability, neuronal loss, and stages behavioral defects (*4-9*). Early transmission studies showed that the mouse-human cross-species seeding barrier was only overcome by the substitution of the human α-syn sequence with specific native mouse residues (*10, 11*). Thus, wild type (WT) mouse PFFs have been assumed to be a “safer” option to recapitulate the main pathologic hallmarks of human synucleinopathies with lower risk of potential infection in humans. More recently, the successful transmission of WT mouse α-syn PFFs into mice that express human α-syn and into human induced pluripotent stem cell (iPSC)-derived neurons proved that the α-syn mouse-human species barrier of transmission is weaker than previously reported (*12*). While no case of work-related transmission of α-syn has been reported to date, these transmission studies prompted the scientific community to reconsider the current biosafety practices when handling mouse α-syn (*13*).

The peripheral transmission of high concentrations of α-syn PFFs or LB-enriched fractions in non-human primates (*14*), rats (*15*), WT mice or mice overexpressing mutant α-syn has been documented (*16-19*). One of the laboratory procedures posing a high risk of accidental human exposure to pathologic α-syn is the sectioning of α-syn-contaminated tissues, particularly brains. Misfolded α-syn aggregates cannot be degraded by the inactivation methods standard in histopathology, such as formalin fixation (*20*). Formic acid is a weak organic acid well-known for its efficiency in prion decontamination (*21, 22*). This acid is often used in α-syn histopathology for antigen retrieval (*23-26*), and thus, has potential to be strategically useful for inactivation of α-syn aggregates in infected tissues. Using real-time quaking-induced conversion (RT-QuIC), we found that formic acid treatment of the brains from DLB patients and PFF-injected mice decreases the seeding activity of α-syn by 6 and 8 logarithms, respectively. Notably, the formic acid treatment did not impact the quality of downstream immunostaining. Together, our studies identify a fast and easy treatment to decrease the seeding activity and, consequently, the risk of accidental exposure to α-syn aggregates in brain tissues.

## RESULTS

### Formic acid treatment decreases the α-syn seeding activity in α-syn-injected mouse brains

Incubating scrapie and Creutzfeldt-Jakob disease (CJD) infected brains with formic acid for an hour drastically reduces the seeding capacity of prions (*21, 22*). To assess the effect of formic acid on α-syn seeding activity, brains from bilateral intrastriatal PFF-injected mice, and the corresponding negative controls (uninjected mouse brains) were cut in six pieces. Frontal and caudal pieces were fixed overnight with 4% paraformaldehyde (PFA), incubated with 96% formic acid (FA) for an hour (right hemisphere) or untreated (left hemisphere controls), and homogenized in RT-QuIC buffer (rostral) (*27*) or immediately immersed in 30% sucrose for immunostaining (caudal) (Fig. 1). The middle brain pieces from both hemispheres were snap-frozen at harvest, remaining unfixed with PFA and untreated with formic acid to confirm that the levels of Triton X-100 insoluble α-syn were similar between brain hemispheres of the same animal (Fig. S1).

**Figure 1.**
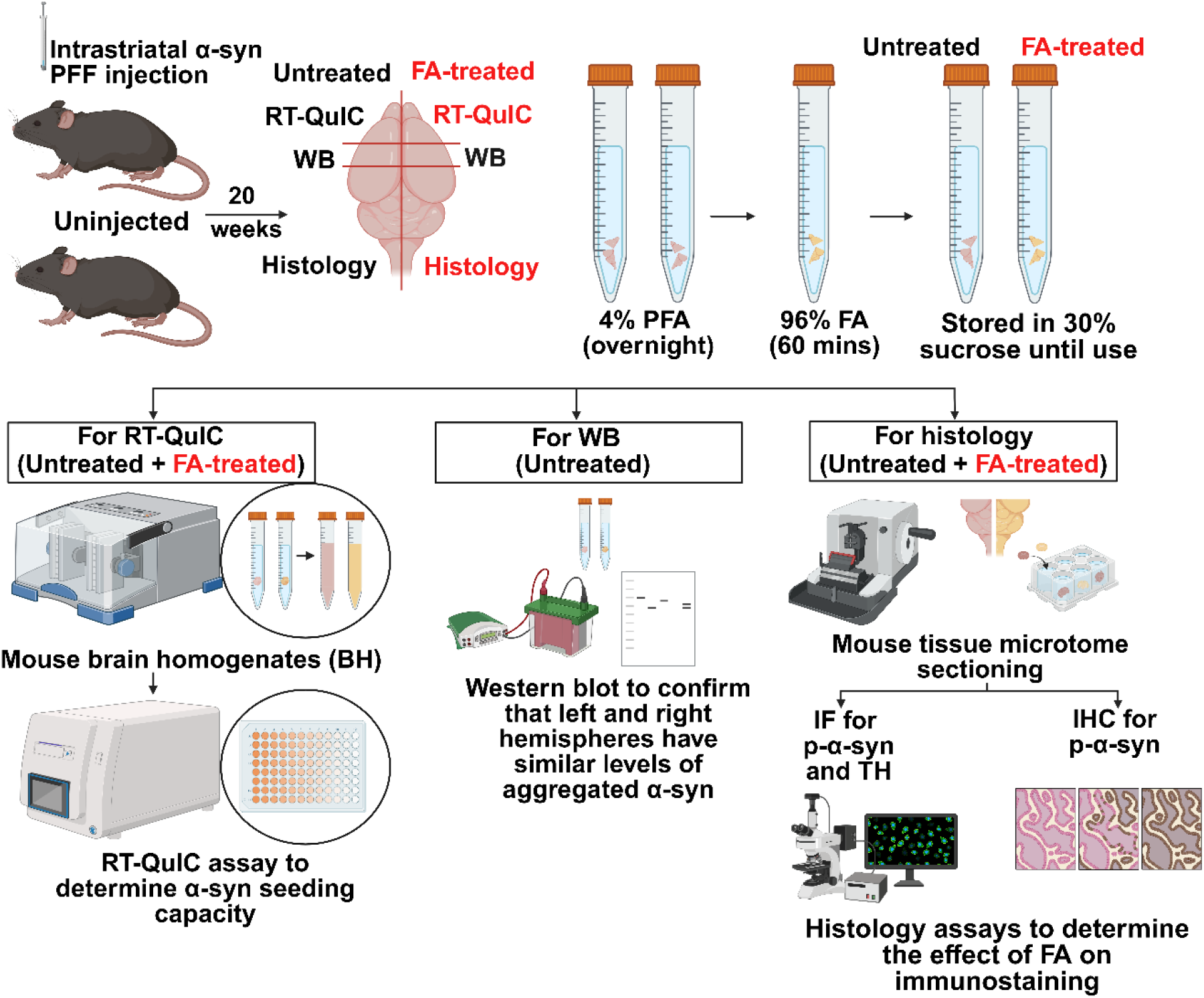
Schematic of experimental procedure. Three-month-old wild-type (WT) mice were bilaterally injected with mouse recombinant α-syn PFFs in the striatum. Twenty weeks post-injection, both PFF-injected and age-matched uninjected control mice were sacrificed and transcardially perfused with PBS. Frontal and caudal regions were fixed overnight at 4°C with 4% paraformaldehyde (PFA), for an hour with formic acid (FA) and stored in 30% sucrose until use. Frontal brain pieces (containing frontal cortex) were processed for RT-QuIC to assess α-syn seeding activity in FA-treated tissues compared to controls. Caudal brain pieces (containing cortex, striatum and substantia nigra) were analyzed by immunofluorescence and immunohistochemistry for phosphorylated α-syn (p-α-syn) and tyrosine hydroxylase (TH) to evaluate the effects of FA treatment on tissue pathology. Middle brain pieces (containing cortex and striatum) were snap-frozen and used for western blot analysis to evaluate the levels of aggregated α-syn among groups. Schematic created with Biorender.com.

The RT-QuIC seeding kinetics of α-syn aggregates were compared between FA-treated and untreated brain homogenates as time-dependent increases in ThT fluorescence intensity above the background threshold (50% increase over the average fluorescence from cycles 3–14 of FA-treated and untreated PFF-injected samples). While RT-QuIC assays seeded with 2 μL of PFF-injected mouse brain homogenates demonstrated strong α-syn seeding activity; RT-QuIC reactions seeded with non-injected brain homogenates showed barely to none seeding activity (Fig. 2A). The PFF-injected untreated samples crossed the fluorescence threshold (9,339.135 RFU) as early as 14.81 ± 0.90 hours and reached maximum fluorescence at 30.75 ± 3.34 hours (Fig. 2B). FA-treated PFF-injected mouse brain homogenates showed significantly reduced ThT fluorescence and crossed the fluorescence threshold at 25.75 ± 4.99 hours (Fig. 2B, Fig. S2). Besides, the maximum fluorescence signal at the end of the assay (33 hours) was markedly lower in FA-treated PFF-injected brain homogenates (FA-treated: 17,131 ± 5,737 RFU vs FA-untreated: 82,430 ± 15,692 RFU) (Fig. 2C), and the lag phase was significantly extended (FA-treated: 25.75 ± 4.99 hours vs FA-untreated: 14.81 ± 0.90 hours) (Fig. 2D), indicating delayed and diminished seeding activity in the FA-treated PFF-injected brain homogenates. Comparative analysis revealed a significantly lower protein aggregation rate (PAR), calculated as the inverse of the lag phase (1/h), in FA-treated PFF-injected samples (FA-treated: 0.0393 ± 0.007 h^-1^ vs FA-untreated: 0.0672 ± 0.004 h^-1^) (Fig. 2E). Collectively, these studies show that one-hour treatment with 96% formic acid impairs α-syn seeding activity in PFF-injected brain homogenates.

**Figure 2.**
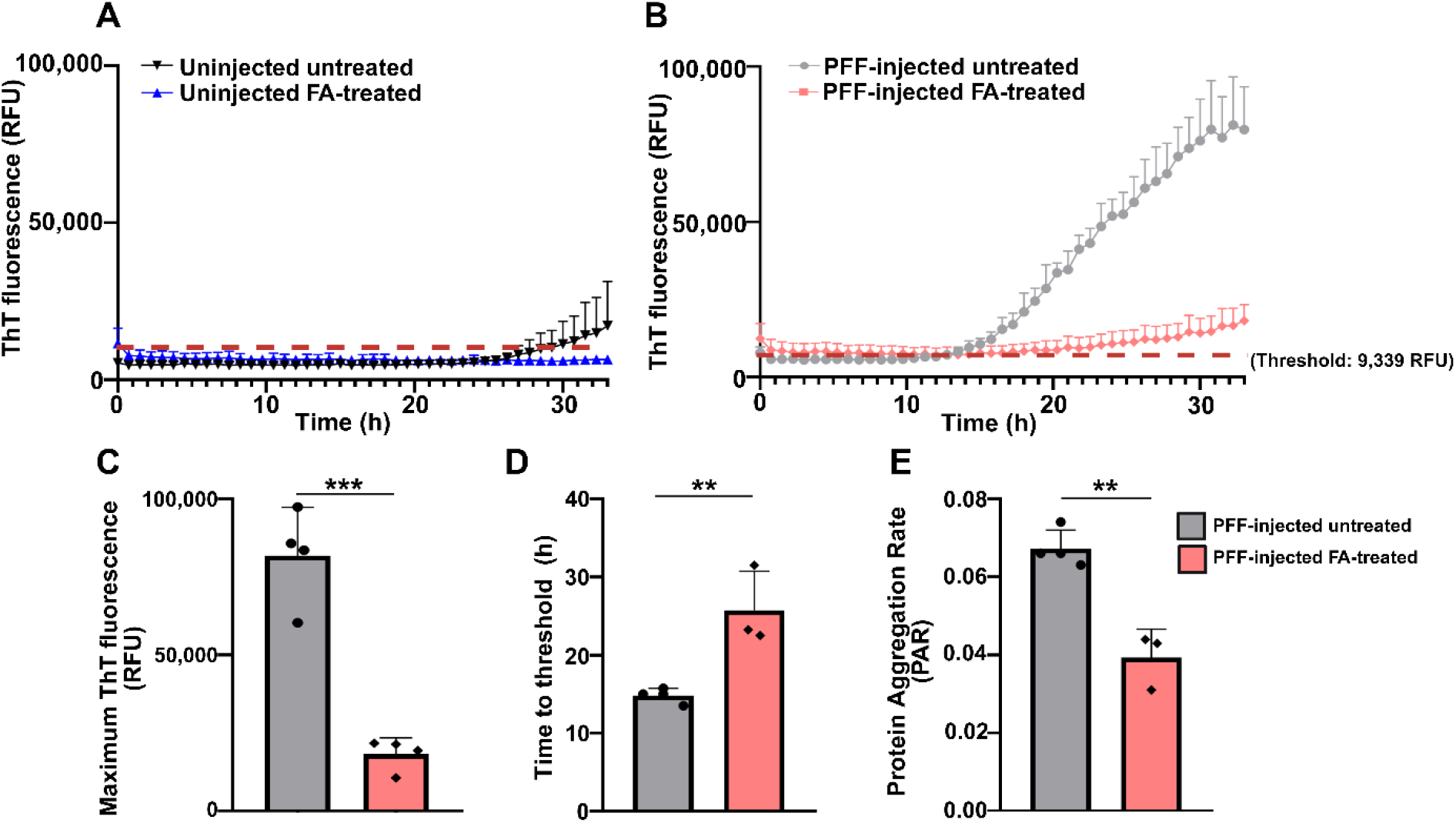
Formic acid-treated PFF-injected mice brain homogenates show a significant decrease in seeding activity. **(A,B)** Summary of RT-QuIC results from seeding the substrate with uninjected (A) and PFF-injected (B) brain homogenates (untreated vs. FA-treated). The threshold (9,339.135 RFU) is indicated on the graphs by a red dotted line. **(C)** Quantification of the maximum fluorescence reveals a significant reduction of ThT fluorescence in the reactions seeded with FA-treated samples. **(D)** The lag phase is significantly increased in reactions seeded with FA-treated samples, indicating a delayed onset of aggregation. Note: Brain homogenate #4 had higher ThT fluorescence than the rest of the FA-treated mice, crossing the threshold at 0 hours, but maintained the slope progression seen in other FA-treated samples (see Fig. S2D). **(E)** The protein aggregation rate (PAR) is significantly reduced following treatment with formic acid. Panels C-E: Unpaired t-test; \*\**P* < 0.01, \*\*\**P* < 0.001.

The α-syn seeding activity of PFF-injected brain homogenates faded with the first logarithmic dilution for untreated and FA-treated samples, which prevented us from applying the RT-QuIC coupled with a modified Spearman-Karber method for relative quantification of proteopathic seeds (*28*). Alternatively, we seeded the RT-QuIC reactions with serial logarithmic dilutions of mouse α-syn PFF of known concentration (5 mg/ml). The α-syn concentration of each PFF-injected mouse brain was calculated from the 50% increase in relative fluorescence units (RFU)–PFF concentration (pg) linear regression line (Fig. S3). After extrapolating the 50% RFU of each untreated and FA-treated brain homogenate from the linear regression (Fig. S3), we found that the approximate estimate of α-syn seeds per brain homogenate volume (ml) for the untreated PFF-injected brain homogenates ranged from 5 to 500 pg of α-syn/ml brain homogenate. In contrast, the estimation of seeds in the FA-treated samples drastically decreased to 1.6×10^−10^ to 5.0×10^−5^ pg/ml brain homogenate. Thus, the formic acid treatment reduced the α-syn seeding dose in PFF-injected mouse brains by approximately 8 logarithms, providing evidence for its utility as a decontamination strategy for α-syn-contaminated mouse tissues in laboratory settings.

### Formic acid treatment decreases the α-syn seeding activity in the brains of DLB patients

Brain tissue samples from the temporal cortex of DLB patients and age-matched healthy control subjects were fixed with 4% PFA overnight, treated with 96% formic acid, and used as seeds for RT-QuIC. While RT-QuIC reactions seeded with brain homogenates from DLB patients showed robust α-syn seeding activity, barely any seeding activity was observed in the reactions seeded with homogenates from healthy control individuals (Fig. 3A). The RT-QuIC reactions seeded with 2 μl of FA-treated DLB brain homogenates crossed the fluorescence threshold (11,974.61 RFU) later than untreated DLB brain homogenates (FA-treated: 31.5 ± 5.25 hours vs FA-untreated: 23.5 ± 3.12 hours) (Fig. 3B, Fig. S4), showed significantly reduced maximum ThT fluorescence at 45 hours (FA-treated: 41,790.17 ± 13,507.33 RFU vs FA-untreated: 171,583.42 ± 11,767.78 RFU), trends towards extended lag phase (FA-treated: 31.5 ± 5.25 hours vs FA-untreated: 23.5 ± 3.12 hours) and lower PAR (FA-treated: 0.0324 ± 0.0059 h^-1^ vs FA-untreated: 0.0430 ± 0.0054 h^-1^) (Fig. 3C-E).

**Figure 3.**
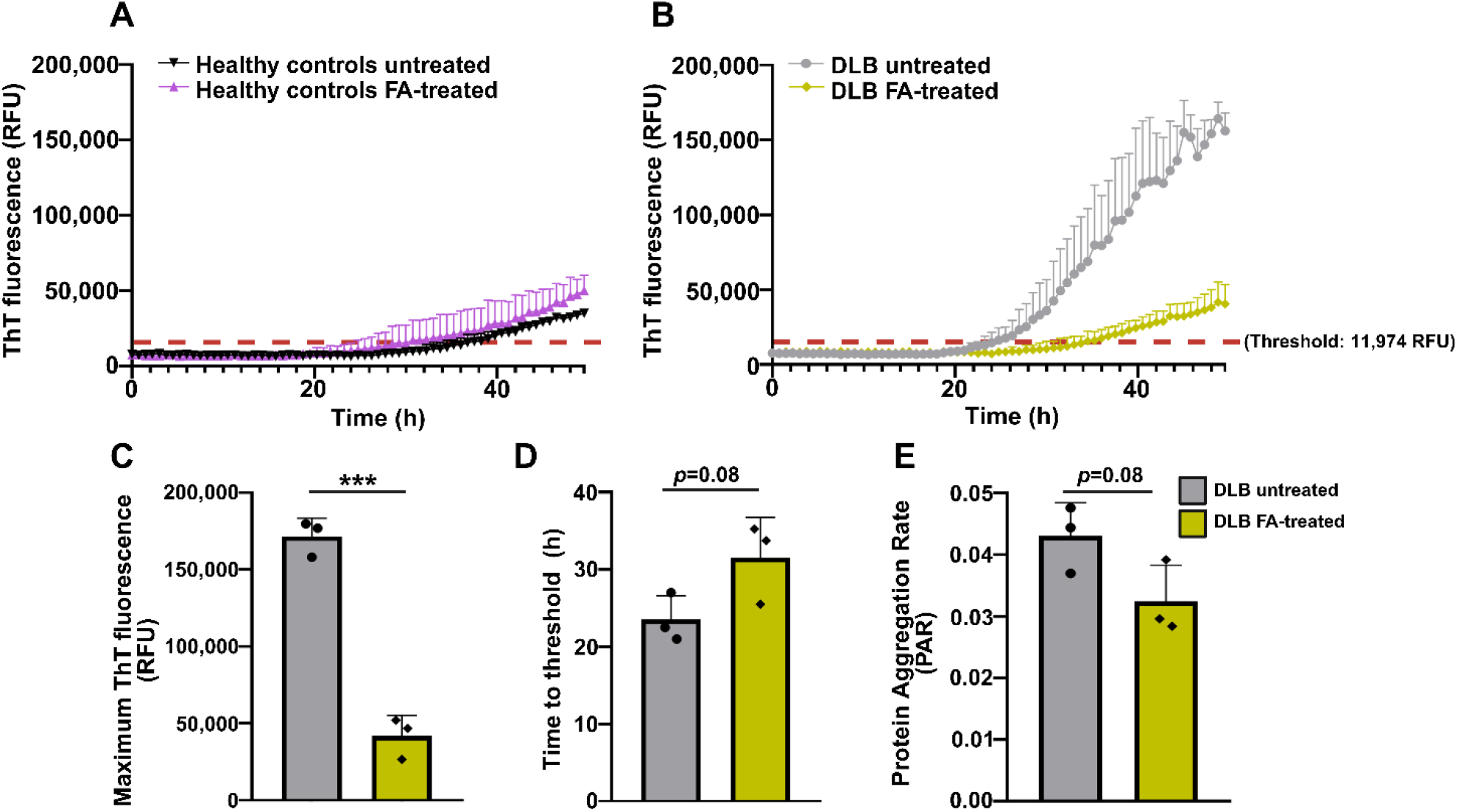
Formic acid-treated human DLB brain homogenates show a marked decrease in seeding activity. **(A,B)** Summary of RT-QuIC results from seeding the substrate with healthy brain controls (A) and DLB brain (B) samples (untreated vs. FA-treated). The threshold (11,974.61 RFU) is indicated on the graphs by a red dotted line. **(C)** Quantification of the maximum fluorescence at the end of the study (45 hours) reveals a significant reduction of ThT fluorescence in the reactions seeded with FA-treated samples. **(D,E)** No statistical differences were observed in the lag phase (D) and the protein aggregation rate (PAR) (E) following formic acid treatment. Panels C-E: Unpaired t-test; \*\*\**P* < 0.001.

To quantify the α-syn seeding activity in DLB brain homogenates, we seeded the RT-QuIC reactions with serial logarithmic dilutions of human α-syn PFF of known concentration (5 mg/ml), and extrapolated α-syn concentrations of each DLB brain homogenate sample using a standard curve (Fig. S5A), as previously described for mouse tissue. We found that the estimate of α-syn seeds per brain homogenate volume for the untreated DLB brain homogenates ranged from 1,584 to 40,738 pg α-syn/ml (Fig. S5B). In contrast, the estimation of seeds in the FA-treated samples was 3.6×10^−3^ to 6.6×10^−1^ pg/ml. Thus, the formic acid treatment reduced the α-syn seeding dose in DLB brains by approximately 6 logarithms, implying it as an effective method for safer human tissue handling.

### Formic acid treatment does not alter the immunostaining of α-syn pathology

Formic acid is a standard epitope retrieval method for α-syn immunohistochemistry (*23*), typically used at concentrations of 88% or lower and with incubation times not exceeding 30 minutes. To determine the impact of one-hour exposure to 96% formic acid on the quality of immunostaining, untreated controls and FA-treated contralateral hemispheres from PFF-injected mice were sectioned and stored in cryoprotectant buffer before immunohistochemistry and immunofluorescence. An initial macroscopic analysis showed that FA-treated brain slices were more rigid and resistant to folding compared to untreated sections, though they were also more brittle. However, immunofluorescence analysis for p-α-syn revealed well-defined LB and LN pathology of both untreated and FA-treated sections (Fig. 4A). The area covered by p-α-syn deposits was similar between both groups (untreated and FA-treated), (Fig. 4B), while the dopaminergic neurons stained with tyrosine hydroxylase (TH) also maintained their characteristic fusiform and elongated appearance in both treatment groups (Fig. 4A). Lewy inclusions were similarly stained using immunohistochemistry (Fig. 4C), confirming that the tissue incubation with formic acid for an hour does not disrupt pathological α-syn detection.

**Figure 4.**
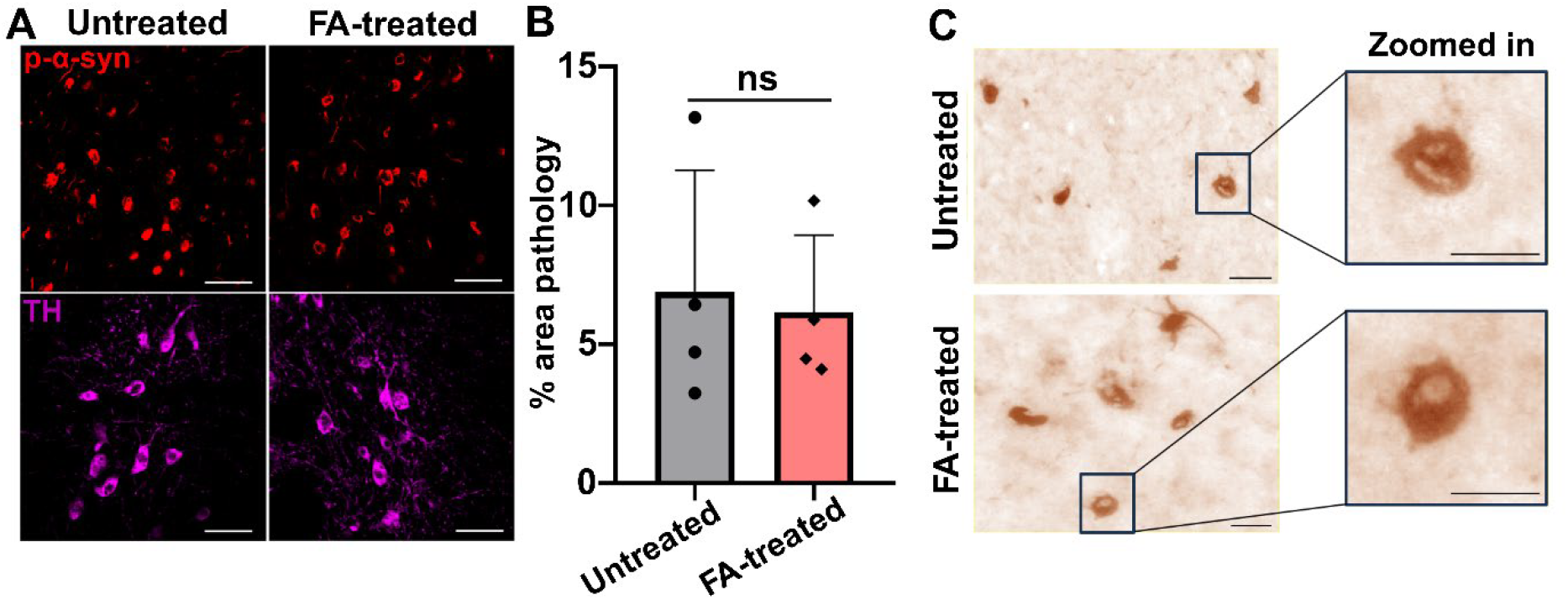
Formic acid treatment does not alter phosphorylated α-syn or dopaminergic neuron staining. **(A)** Immunofluorescence images showing phosphorylated α-syn (p-α-syn) pathology in the amygdala and dopaminergic neurons stained with tyrosine hydroxylase (TH) in the substantia nigra pars compacta of PFF-injected untreated and FA-treated mouse brain samples. No observable differences were noted between groups. Scale bar = 50 μm. **(B)** Quantification of p-α-syn pathology in the cortex is represented as % area covered by pathology. No significant (ns) difference was observed between the untreated and FA-treated groups. Unpaired t-test. **(C)** Immunohistochemistry images of p-α-syn in PFF-injected untreated, and FA-treated animals. Images show Lewy bodies in the cortex from both groups, with zoomed-in insets highlighting the inclusion structure. No notable differences in Lewy body stain or appearance were observed between the groups. Scale bar = 20 μm.

## DISCUSSION

The recently reported high seeding capacity of mouse α-syn fibrils in humanized mouse brain and human-iPSCs (*12*) has prompted a reassessment of the potential risk from manipulating α-syn-contaminated tissues (*13*). Implementing safety measures to work with α-syn fibrils and contaminated tissues has been significantly hampered by the scarce information on the reagents or conditions that affect the conformational stability and seeding capacity of α-syn fibrils. Our RT-QuIC studies show that an hour immersion with 96% formic acid drastically reduces α-syn seeding activity up to 6 logarithms in DLB brains and up to 8 logarithms in PFF-injected mouse brains. Moreover, we found that the seeding dose in the DLB and PFF-injected mouse brains reached 10^−3^ and 10^−10^ pg/ml, which are drastically lower than the 10 to 50 μg of mouse PFFs needed to induce α-syn pathology in mice peripherally (*18, 29*). Since the quality of histopathological analysis is not significantly affected by the treatment with this acid, adding a 1-hour formic acid incubation step to the routine formaldehyde fixation of brain tissue appears to be the best procedure yet devised for minimizing the potential exposure to pathological α-syn.

Formaldehyde fixation does not inactivate α-syn seeds (*30*). Hypochlorous acid abolished the seeding activity of recombinant α-syn fibrils (*31*), whereas autoclaving at 121°C for 20 minutes removes one to two logarithms of α-syn seeding activity in the brains of DLB patients (*32*). However, the suitability of these inactivating techniques for histopathology analysis is unknown. Formic acid has higher protein solvent activity than most common organic solvents including glycerol and DMSO, solubilizing the protein polypeptide chain through protonation, destabilization of hydrogen bonds, and hydrophobic residue interactions (*33*). We found that a one-hour incubation with formic acid reduced the seeding activity of α-syn in PFF-injected mouse brains approximately by 8 logarithms and up to 6 logarithms in DLB brains. These results are similar to those previously observed when treating prion-contaminated human and mice tissues with high concentrations of formic acid. Using mouse bioassay, Brown et al. found 8 logarithms loss of infectious units (obtained from the mean lethal dose) in prion-infected hamster brains treated with formic acid (*21*); whereas CJD-infected brains showed a loss of 7 orders of magnitude in the infectious units (*22*). In this study, α-syn seeding activity was measured by RT-QuIC, a faster but similarly sensitive technique compared to mouse bioassay (*28*). Despite the differences in the original seeding loads in the prion-and α-syn-contaminated tissues, the assays and conditions used to quantify seeding activity or the conformational stability of the seeds, others’ and our studies demonstrate that formic acid is highly effective at inactivating prion and α-syn seeds.

How does formic acid reduce the seeding activity of pathological α-syn? The denaturing capacity of formic acid was extensively investigated for the functional amyloid protein FapC using near-ultraviolet circular dichroism, a highly sensitive technique to correct folding of the protein (*34*). These studies showed that formic acid destabilizes the FapC protein by altering its secondary structure, disrupting the formation of stable amyloid fibrils. Notably, the authors found that the denaturing effect of formic acid directly correlates with the number of imperfect repeats in the FapC protein, a motif of repeating amino acid residues needed for fibril aggregation and resistance to fragmentation (*34*). The N-terminal and non-amyloidal component (NAC) regions of mouse and human α-syn contain seven imperfect 11 animo acid repeats (XKTKEGVXXXX) that include a highly conserved KTKEGV hexameric motif. Mounting evidence indicates that these imperfect repeat domains are key for modulating the kinetics of fibrillation and the morphology of fibrils (*35-37*), and mutations in the imperfect repeats deeply affect the kinetics of fibril formation and the morphology of fibrils. Exposure to formic acid leads to protonation of glutamate residues (E–COOH), disrupting these stabilizing interactions and the β-sheet structure (*34*). This might result in the generation of seeding-incompetent fibrils, as illustrated by RT-QuIC. The N-terminus of the prion protein contains a four to five octapeptide repeat region (octarepeats). An increase of the octarepeat numbers to six or more or a decrease of the octarepeat number to three is linked to genetic prion diseases with heterogeneous phenotypes in humans (*38, 39*). Thus, we suggest a common mechanism of protein seeding capacity inactivation based on the ability of formic acid to interact with the imperfect repeats amino acids and alter the protein conformational stability.

As our knowledge of the α-syn fibrils structure grows, more methods will emerge to decrease the seeding activity of pathological α-syn. The inactivation procedure described here is fast, straightforward, and provides the greatest reduction in α-syn seeding activity reported to date without significantly affecting the quality of the immunostaining analysis or the tissue structure. Thus, we propose incubating α-syn-contaminated tissues for 1 hour with 96% formic acid to minimize the potential exposure of scientists, neuropathologists, and other workers to pathological α-syn while manipulating contaminated tissues. Lastly, since the tau protein contains three or four 31-32-residue imperfect repeats that are key for in vitro fibril formation (*40-42*), formic acid treatment might also decrease tau seeding activity in the brains of Alzheimer’s disease and frontotemporal dementia patients as well as animal models of tauopathies.

## MATERIALS AND METHODS

### Study design

The objective of this study was to assess the effectiveness of formic acid treatment for α-syn inactivation in tissue. To this end, sections from PFF-injected mice (N= 4) or the temporal cortex of DLB patients (N= 3) were either treated for one hour with formic acid or not, homogenized, and then used as seed in RT-QuIC. This rapid, quantitative, and high-throughput assay for in vitro replication of protein aggregates allowed us to define the impact of formic acid on the seeding capacity of α-syn in each sample. All the homogenates were tested in technical quadruplicates. To determine the effect of the formic acid treatment on tissue quality, untreated and FA-treated sections from PFF-injected mice (N= 4) were used for immunofluorescence and immunohistochemistry against p-α-syn and dopaminergic neurons’ marker tyrosine hydroxylase (TH). Two researchers blinded to the tissue treatment scored the morphology and distribution of the α-syn inclusions in mouse brains.

Human autopsy brain tissue samples were provided by the Emory University Goizueta Alzheimer’s Disease Research Center (GADRC) brain bank. Details on these samples are provided below. All animal studies were approved by the Institutional Animal Care and Use Committee at University of Alabama at Birmingham (IACUC protocol #22603). Protocols were performed in strict accordance with good animal practices, as described in the Guide for the Use and Care of Laboratory Animals published by the National Institutes of Health. C57BL/6 J mice were maintained on a 12 hours light/dark cycle with unrestricted access to food and water.

### Preparation of recombinant α-syn preformed fibrils

α-Syn preformed fibrils (PFFs) were generated from recombinant monomeric α-syn following a previously published protocol (*43*). Briefly, α-syn monomers were expressed in *E. coli*, purified using ion exchange and size exclusion chromatography, and verified for purity by SDS-PAGE before fibril assembly. α-Syn monomers were incubated with 150 mM KCl and 50 mM Tris HCl at 37°C and agitated for 7 days at 1,000 rpm. Fibrils were later isolated from the monomers by centrifugation for 10 minutes at 13,200 rpm, their protein concentration was determined by spectrophotometer, and a buffer consisting of 50 mM Tris-HCl and 150 mM KCl was used to bring the concentration to 5 mg/ml. PFFs were stored at −80°C until use.

Immediately before injections, PFFs were sonicated using a Qsonica sonicator with circulating water at 15°C. Samples at 22 μL were placed in a 1.5 ml sonicator tube (Active Motif, NC0869649) and sonicated for 15 minutes (amplitude 35; pulse on 3 seconds; pulse off 2 seconds). Fragmentation of PFFs between 20 and 30 nm fragments was confirmed using dynamic light scattering (Wyatt Technology).

### Intrastriatal injection of recombinant α-syn fibrils and tissue sampling

At three months of age, mice (N= 4) were deeply anesthetized with vaporized isoflurane and secured on a stereotaxic frame. Mice were then bilaterally injected with 2 μl (5 mg/ml) of sonicated α-syn fibrils (10 μg total protein per mouse) into the dorsolateral striatum (+1.0 mm AP and ±2.0 mm ML from bregma; -2.6 mm DV from dura). Fibrils were injected at a constant rate of 0.5 μl/min. Once the injection was complete, the needle remained in place for 4 minutes before being gradually withdrawn.

At 20 weeks post-injection, PFF-injected mice and age-matched non-injected controls were anesthetized with isoflurane and perfused transcardially with 1x phosphate-buffered saline (PBS). During necropsy, the brain was sectioned as illustrated in Fig. 1, with the frontal and caudal regions immediately fixed in 4% paraformaldehyde at 4°C. After 24 hours, only the right frontal and caudal regions were treated with 96% formic acid for one hour, then submerged in cryoprotectant (30% sucrose in PBS) until use. Caudal sections from both hemispheres were cut at 40 μm thickness using a Leica SM 2010 R freezing microtome. Serial sections were collected in 6-well plates, ensuring each well contained slices spaced 240 μm apart, representing the entire brain region. Brain sections were stored in a cryoprotectant buffer of 50% glycerol, 0.01% sodium azide in Tris-buffered saline (TBS) until use. The middle sections were immediately frozen in dry ice for immunoblotting analysis.

### Immunostaining

For immunofluorescence, sections were washed with cold 1x TBS and subjected to antigen retrieval by incubating in preheated 10 mM sodium citrate buffer (pH 9.0) at 80°C for 15 minutes. Following retrieval, sections were washed with cold 1x TBS and blocked with blocking buffer (10% normal goat serum (NGS), 0.1% Triton-X-100 in 1x TBS) for one hour. Primary antibody incubation was performed using a solution containing Rabbit Anti α-syn (Ser129) antibody (Abcam, ab51253; 1:500) and Chicken Anti-tyrosine hydroxylase antibody (Abcam, ab76442; 1:500) in 10% NGS in 1x TBS for 48 hours at 4°C. Sections were washed with cold 1x TBS and incubated with secondary antibodies: Cy3 AffiniPure Goat Anti-rabbit IgG (H+L) (Jackson ImmunoResearch Laboratories, 111-165-144; 1:500) and Cy5 Alexa Fluor 647 Goat Anti-Chicken IgY (H+L) (Thermo Fisher Scientific, A21449; 1:500) in 1x TBS with 10% NGS for 2 hours at 4°C. Following the final wash in cold 1x TBS, sections were mounted onto positively charged slides, allowed to dry, and coverslipped with ProlongGold.

Pathology was analyzed by two researchers who were blinded to the experimental conditions. Sections were imaged using a VS200 slide scanner (Olympus) at 20X magnification, visualized with QuPath software, and analyzed with ImageJ software. The region of interest (ROI) spanning the cortex was manually marked, and the percentage area covered by inclusions was measured by setting a consistent threshold. Representative images were imaged on a Nikon C2 confocal microscope at 40X and visualized using Nikon Elements software.

For immunohistochemistry, free-floating brain sections were washed in cold 1x TBS and incubated in 3% hydrogen peroxide for 20 minutes at room temperature (RT) to quench endogenous peroxidase activity, followed by rinsing in distilled water. Antigen retrieval was performed by incubating sections in pre-heated sodium citrate buffer (pH 9.0) at 80°C for 15 minutes in a water bath. Sections were then blocked in a solution containing 5% NGS and 0.05% Triton X-100 in 1x TBS for 30 minutes at 4°C with gentle agitation. Following blocking, tissues were incubated for 48 hours at 4°C in primary antibody solution containing Rabbit Anti α-syn (Ser129) antibody (Abcam, ab51253; 1:4,000) diluted in 5% NGS and 0.05% Triton X-100 in TBS. To prevent tissue adhesion, sections were incubated individually in separate wells of a 24-well plate. After primary incubation, sections were washed and incubated in secondary antibody solution containing Peroxidase AffiniPure Goat Anti-Rabbit IgG (H+L) (Jackson ImmunoResearch, 111-035-003; 1:1,000) for two hours at RT. Immunoreactivity was visualized using the ImmPACT DAB substrate kit (Vector Laboratories, SK-4105); sections were developed for ∼1.5 minutes, and the reaction was terminated by rinsing in distilled water. Finally, sections were washed, mounted onto glass slides in 1x TBS, air-dried, and coverslipped using Permount for imaging. Slides were images using VS200 slide scanner (Olympus) at 40X and visualized using QuPath software.

### Western blot

Mouse brains were homogenized and sonicated in lysis buffer (50 mM Tris-HCL pH 7.4, 175 mM NaCl, and 5 mM EDTA pH 8.0) with 1% Triton X-100, incubated on ice for 30 minutes, and spun for 60 minutes at 15,000×g to extract Triton X-100-soluble proteins. Pellets were then resuspended in lysis buffer containing 2% SDS and sonicated to extract Triton X-100-insoluble proteins. Samples were immunoblotted for α-syn following proteinase K digestion of the SDS-soluble fraction. Protein concentrations were assessed by BCA assay (Thermo Fisher Scientific). Tissue lysates were boiled in 4x DTT sample loading buffer (0.25M Tris-HCl pH 6.8, 8% SDS, 200 mM DTT, 30% glycerol, Bromophenol Blue). Equal amounts of protein (20 μg) were loaded per well, resolved on 4-20% SDS-polyacrylamide gels, and transferred to nitrocellulose membranes. After blocking in 5% nonfat dry milk in TBST (25 mM Tris-HCl, pH 7.6, 137 mM NaCl, 0.05% Tween 20), membranes were incubated overnight at 4°C in primary antibody solution containing Mouse Anti α-syn (BD Transduction Laboratory, 610787; 1:1,000) and then in HRP-conjugated Goat Anti-mouse or Anti-rabbit secondary antibodies (Jackson ImmunoResearch; 115-035-003; 1:2,000 or Jackson ImmunoResearch; 111-035-003; 1:2,000) for one hour at RT. After washing with TBST, blots were developed with enhanced chemiluminescence (GE Healthcare) and imaged with a ChemiDoc XRS+ (Bio-Rad Laboratories).

### Real-time quaking-induced conversion assay

The RT-QuIC assay for mouse and human α-syn was performed per published work (*27*), with modifications. Briefly, RT-QuIC reactions were performed in Nunc black 96-well plates with optical flat bottom (Thermo Fisher, Waltham, MA, USA). Each well was preloaded with six 0.8 mm low-binding silica beads (OPS Diagnostics, Lebanon, NJ, USA). Lyophilized recombinant mouse or human α-syn (rPeptide) was reconstituted in HPLC-grade water to 1 mg/ml and filtered through Amicon 100 kDa filters (Millipore, Burlington, MA, USA) by centrifugation for 10 minutes at 4°C. To measure α-syn RT-QuIC seeding activity, 2 μl of diluted tissue homogenate was added to individual wells containing 98 μl of RT-QuIC reaction mixture composed of 40 mM NaPO4 pH 8.0, 170 mM NaCl, 20 μM ThT, and 0.1 mg/ml recombinant mouse or human α-syn. The plates were sealed with Nunc clear sealing film (Thermo Fisher, Waltham, MA, USA) and incubated at 38°C in a BMG FLUOstar Omega plate reader (BMG Labtech, Cary, NC, USA) with cycles of 1 minute shaking (400 rpm, double orbital) and 15 minutes rest throughout the assay. ThT fluorescence (450 nm excitation and 480 nm emission; bottom read) was recorded every 45 minutes for the duration of assay.

### Dementia with Lewy bodies patients

Brain tissue samples from three patients with dementia with Lewy bodies (DLB) and two healthy controls (Table 1) were used to investigate how formic acid treatment affects the seeding activity of α-syn in tissue. The DLB samples originated from patients enrolled in the Emory University GADRC research program and met neuropathologic criteria (*44, 45*) for the diagnosis of neocortical Lewy body disease; as is commonly seen in DLB, these cases also exhibited Alzheimer’s disease pathology. The Lewy pathology was scored using the Braak standard scoring for neurofibrillary pathology (*46*). The concurrent Alzheimer’s pathology was measured using the ABC scoring per NIA-AA guidelines (*47*), the CERAD scoring for neuritic plaque pathology (*48*), and the Thal scoring for amyloid-beta deposition (*49*). The demographic information, and pathological scoring can be found in Table 1. The mean age of DLB patients at disease onset was 64 ± 14 years (mean ± SD), and the duration of clinical neurologic signs ranged from 2 to 6 years (5 ± 2 years).

**Table 1.** Description of the human cases used for this study.

| Case # | Gender | Age at onset<br>(years) | Age at death<br>(years) | APOE<br>alleles | Braak<br>Stage | ABC | CERAD | Thal | PMI*<br>(hours) |
| --- | --- | --- | --- | --- | --- | --- | --- | --- | --- |
| DLB -1 | female | 64 | 70 | E3/4 | V | 3 | 3 | 5 | 7.5 |
| DLB-2 | female | 50 | 52 | E3/4 | VI | 3 | 3 | 5 | 12 |
| DLB-3 | female | 77 | 83 | E3/3 | V | 3 | 3 | 5 | 5.5 |
| Control-1 | male | - | 53 | E4/4 | II | 0 | 0 | 0 | 6.5 |
| Control-2 | female | - | 78 | E3/3 | II | 0 | 0 | 0 | 11.5 |

\* Post-mortem interval: time between death and when a body is examined.

### Statistical analysis

Statistical analyses were performed using GraphPad Prism 8 software (GraphPad Software, San Diego, CA). Data are presented as mean ± standard deviation (SD). Between-group comparisons were conducted using Student’s t-test for two-group comparisons. For multiple comparisons, one-way analysis of variance (ANOVA) was performed, followed by Tukey’s post-hoc test to identify specific group differences. Statistical significance was defined as a *P*-value < 0.05.

## Funding

National Institutes of Health grant R00AG061251 (PA-C)

American Parkinson’s Disease Association (PA-C)

National Institutes of Health grant R01AG082352 and R01NS118760 (SC)

National Institutes of Health grant R01AG081433 (LV-D)

National Institutes of Health grant P30AG066511 (MG)

## Author contributions

Conceptualization: PA-C, SC, SD

Methodology: PA-C, SC, LV-D, SD, GD, PD, RB, KS, MSG

Investigation: SD, GD, PD, RB, AS, JH Visualization: SD, GD, AS

Funding acquisition: PA-C, SC, LV-D, MG Project administration: PA-C Supervision: PA-C

Writing – original draft: PA-C, SD

Writing – review & editing: PA-C, SD, SC, LV-D, GD, PD, RB, AS, JH, KS, MSG, MG

## Data and materials availability

All the RT-QuIC raw data, western blots, and histology images are available in the main text or the supplementary materials.

