## Supplemental Material for "Formic acid impairs α-synuclein seeding activity in human and mouse brains"

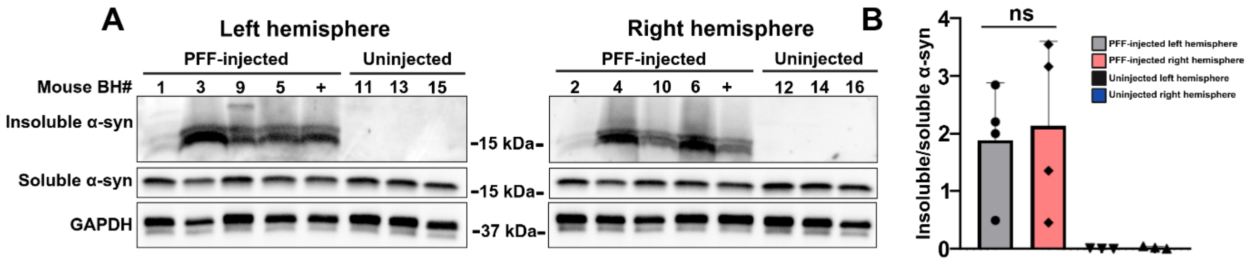


**Sup. Figure 1. Total α-syn levels are similar in the left and right brain hemispheres of each mouse. (A)** Representative western blots showing total levels of Triton X-100 soluble and insoluble α-syn levels in the left (odd numbers) and right (consecutive even numbers) hemispheres from PFF-injected (N= 4) and uninjected (N= 3) mice. A previously analyzed sample from a PFF-injected mouse was used as positive control (+). GAPDH was used as loading control. **(B)** Densitometric analysis of Triton X-100 soluble and insoluble α-syn levels, normalized to GAPDH, shows no significant differences (ns) in the levels of α-syn between left and right hemispheres. Unpaired t-test. Data is presented as mean ± SD.

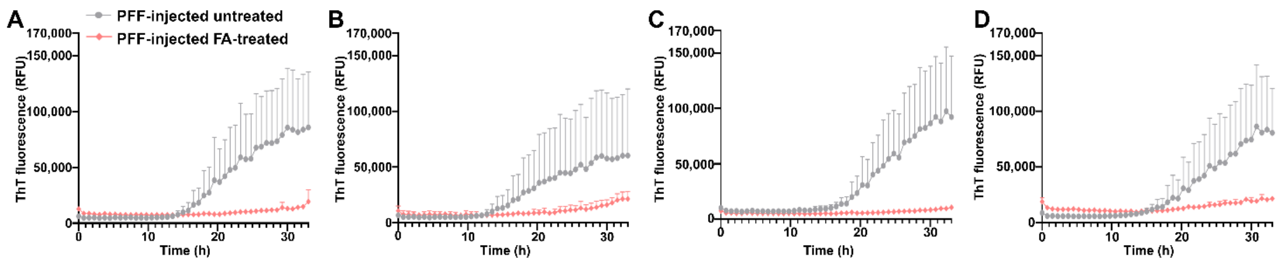


**Sup. Figure 2. Formic acid treatment decreases α-syn seeding activity in PFF-injected mice. (A-D)** Individual RT-QuIC curves from four PFF-injected samples show decreased fluorescence (RFU) in reactions seeded with FA-treated groups.

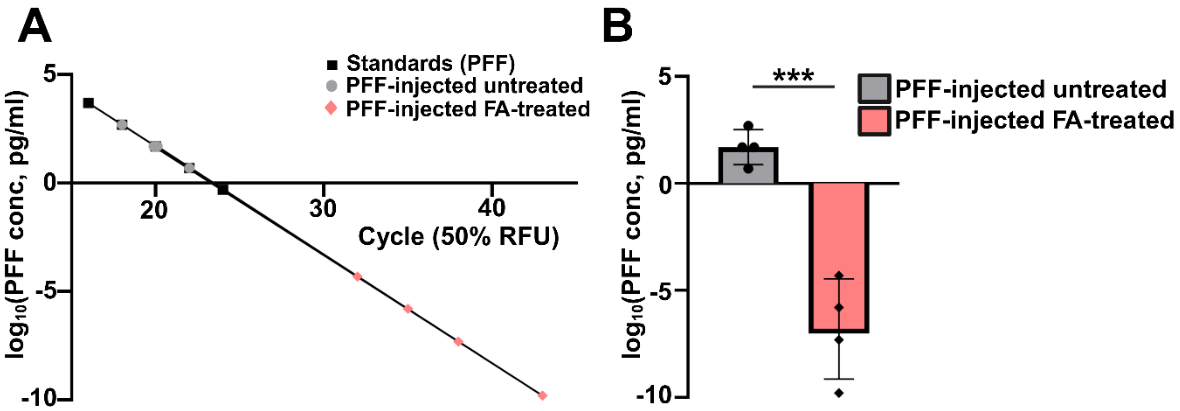


**Sup. Figure 3. Formic acid treatment drastically reduces α-syn seed levels in PFF-injected mice. (A)** Graph shows standard curve generated by linear regression ($R^2$=1.00) using serial dilutions of recombinant mouse α-syn PFFs (5 mg/ml). Estimated seeding concentrations for untreated and FA-treated mouse brain homogenates were extrapolated based on the cycle at which samples reached 50% of maximum ThT fluorescence. **(B)** Graph shows seeding concentrations ($\log_{10}$ pg/ml) extrapolated for untreated and FA-treated mouse brain homogenates. Untreated samples show high seeding activity ranging from 5 to 500 pg α-syn/ml brain tissue homogenate, while FA-treated samples showed drastically reduced levels ($1.6\times10^{-10}$ to $5.0\times10^{-5}$ pg/ml). Unpaired t-test; ****P* < 0.001.

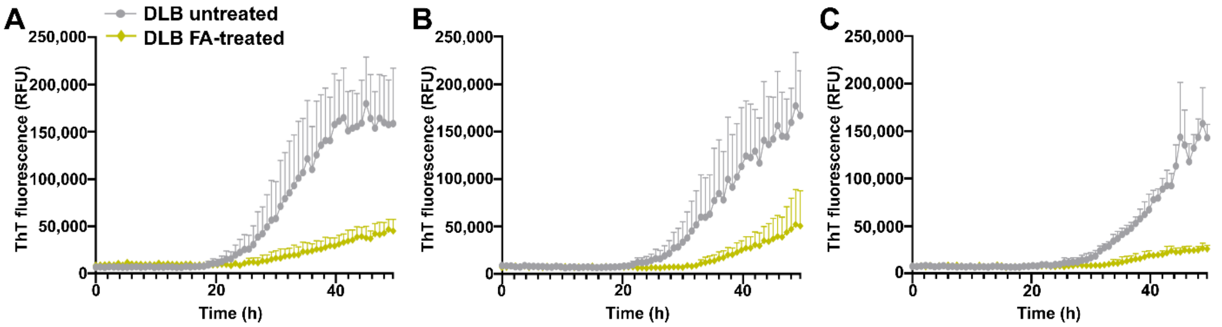


**Sup. Figure 4. Formic acid treatment decreases α-syn seeding activity in human DLB brain homogenates. (A-C)** Individual RT-QuIC curves from three human DLB samples show decreased fluorescence (RFU) in reactions seeded with FA-treated groups.

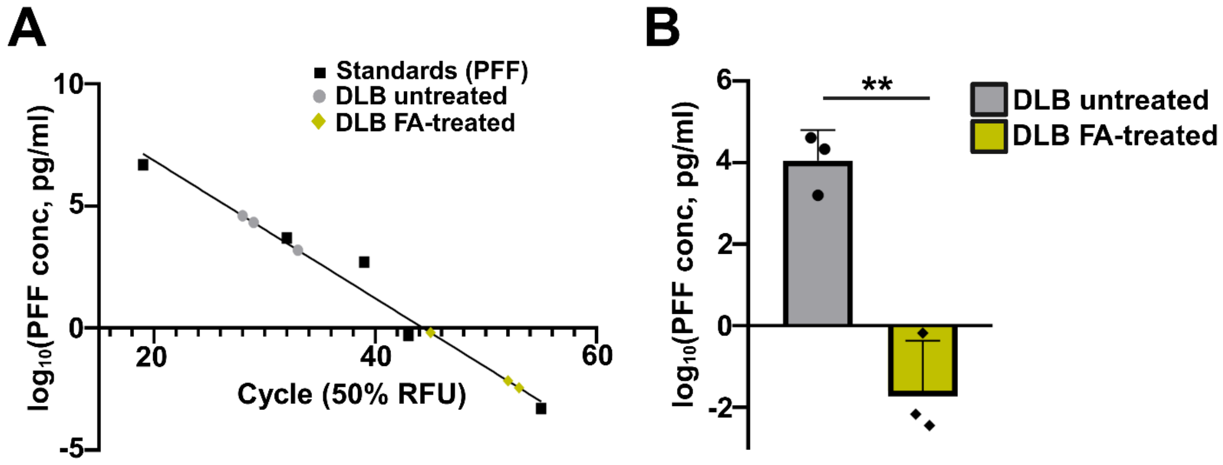


**Sup. Figure 5. Formic acid treatment decreases α-syn seed levels in DLB patient tissue. (A)** Graph shows the standard curve generated by linear regression ($R^2$=0.98) using serial dilutions of recombinant human α-syn PFF (5 mg/ml). Estimated seeding concentrations for untreated and FA-treated DLB brain homogenates were extrapolated based on the cycle at which samples reached 50% of maximum ThT fluorescence. **(B)** Graph shows seeding concentrations ($log_{10}$ pg/ml) extrapolated for untreated and FA-treated DLB human brain homogenates. Untreated samples show increased seeding activity ranging from approximately 1,584 to 40,738 pg α-syn/ml brain tissue homogenate, while FA-treated samples showed decreased levels ($3.6\times10^{-3}$ to $6.6\times10^{-1}$ pg/ml). Unpaired t-test; $**P < 0.01$.
